# Allelic Transcription Factor binding shape transcriptional kinetics observed with scRNA-seq in human cell lines

**DOI:** 10.1101/2022.09.21.508743

**Authors:** Bowen Jin, Hao Feng, William S. Bush

## Abstract

Gene expression from bulk RNA-seq studies is an average measurement between two chromosomes and across cell populations. Both allelic and cell-to-cell heterogeneity in gene expression results from promoter bursting patterns that repeatedly alternate between an activated and inactivated state. Increased cell-to-cell heterogeneity in gene expression has been associated with aging and stem cell pluripotency. However, studies of bursting kinetics and their molecular mechanism are relatively limited in human cells compared to other species due to laborious single-molecule experiments. Here, we systematically investigate the regulatory effect of genetic variants and transcription factor (TF) binding on transcriptional kinetics at the single chromosome level with GM12878. We found that the transcription initiation rate and burst frequency correlate most with eQTL effect sizes among transcriptional kinetics, which suggests that eQTLs affect average gene expression mainly through altering burst kinetics. We further found that ∼90% of the variance of burst frequency can be explained by TF occupancy in phase with the core promoter. We identified and replicated several examples where eQTL or GWAS catalog loci perturb TF binding affinity and are consequently associated with the change of burst kinetics.

## Introduction

Transcriptional bursts are a major contributor to the stochasticity in gene expression in eukaryotic cells (Kærn et al., 2005), which reflects the complexity and hierarchical nature of transcriptional regulation. Transcriptional bursts can introduce significant cell-to-cell transcriptional variability over a cell population and may propagate to phenotypic variability at the tissue or organism level. For example, aging has been shown to increase cell-to-cell transcriptional variability (Bahar et al., 2006; Martinez-Jimenez et al., 2017). Transcriptional bursts can also produce initial heterogeneity in the homogenous cell population, which is further selected and propagated to generate patterns of cell-type specific expression (Chang et al., 2008; Ohnishi et al., 2014; Kumar et al., 2014; Gulati et al., 2020). Hence, understanding more about transcriptional bursts will provide us with detailed molecular mechanisms of transcription and robust ways to characterize different cell types.

Existing studies of bursting expression are mostly single-molecule studies targeted to genes on a single chromosome. Single-molecule techniques like MS2 tagging and Cas-derived systems allow real-time observation of burst kinetics in the context of perturbations to regulatory mechanisms (Singh et al., 2010; Dar et al., 2012; Fukaya et al., 2016; Nicolas et al., 2018; Li et al., 2018; Bartman et al., 2019; Dobrinic et al., 2021). To enable a transcriptome-wide and unperturbed characterization of bursting expression, several other studies explore the potential of using single-cell RNA-seq (scRNA-seq) (Faure et al., 2017; Larsson et al., 2019; Gupta et al., 2022). scRNA-seq has the unique advantage of allowing a snapshot of each cell within the population, capturing the distribution of gene expression over time resulting from the transcriptional bursts. However, scRNA-seq still measures the mixed expression from two chromosomes. We can obtain allelic-level scRNA-seq expression profiles with additional genotype information for allelic-specific transcript mapping. In comparison, existing methods that can explore ASE with single-molecule techniques require laborious genetic manipulation in selected model organisms. Consequently, measurements of bursting expression and its regulatory mechanisms are scattered among various cell lines and experimental systems. In addition, there is a lack of comprehensive, unperturbed, and transcriptome-wide study of human bursting gene expression because it requires comprehensive haplotype profiles, which are only available by familial inference, long-read sequencing, or haplotype inference.

For this study, we designed a pipeline with multiple filtering and validation procedures to characterize bursting kinetics from scRNA-seq data paired with phased genotype profiles. Several studies have developed similar approaches by mapping transcript reads with heterogeneous single nucleotide variants (SNVs) (Borel et al., 2015; Prashant et al., 2021) and fitting the allele level expression profile with a two-state model (Kim & Marioni, 2013; Vu et al., 2016; Jiang et al., 2017; Larsson et al., 2019). These approaches are for full-length scRNA-seq, while our approach is designed for 3’ gene expression with unique molecular identifiers (UMIs), which constitute the bulk of published scRNA-seq datasets.

We used GM12878, the cell line with the most abundant CHiP-seq data available from the ENCODE project, to systematically investigate the molecular regulation of burst gene expression at single chromosome resolution. We derived the allelic occupancy profile of 142 transcription factors (TFs), 8 histone markers, and allelic chromatin accessibility profiles from ATAC-seq data on GM12878. We estimated the variance of transcriptional kinetics explained by the TF binding within the core promoter region and the distal enhancer region in phase with the promoter, respectively. We also investigated how individual TF alters transcriptional kinetics across the genome. Finally, we investigated if eQTLs are associated with the change of transcriptional kinetics due to allele-specific binding (ASB) of TF.

## Results

### Transcriptomic mapping of transcriptional kinetics in lymphoblastoid cell lines of European and African ancestry

We estimated the transcriptional kinetics for GM12878 and GM18502, which are lymphoblastoid cell lines (LCLs) from two females of European descent and African descent, respectively. In detail, we obtained allele-level expression profiles with scRNA-seq data and phased genotype profiles fitted with a beta-binomial distribution (see Methods). While the genotype phase is established for both cell lines, the maternal and paternal origin of chromosomes is only available for GM12878; therefore, for simplicity, we assign one allele to Haplotype1 and the counterpart to Haplotype2 for the context of all analyses.

GM12878 and GM18502 have shown highly consistent ASE patterns (Supplemental Figure S1) and allelic transcriptional kinetics (Figure 2D). Their transcription initiation rate (k+) and transcription termination rate (k-) are not independent of each other and occupy similar restricted phase space (Figure 2A). The average active fraction 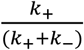 is bound between 0 to 0.6 for both cell lines, and most observations are smaller than 0.5 (Figure 2B). The burst size and burst frequency, empirically independent bursting kinetics derived from k+, k-, and r, (Figure 2C) show similar distributions and boundaries to those derived from mouse primary fibroblasts (Larsson et al., 2019) (Supplemental Figure S2). In conclusion, GM12878 and GM18502 show consistent allelic expression and transcriptional kinetics for the vast majority of genes they express.

**Figure 1.**
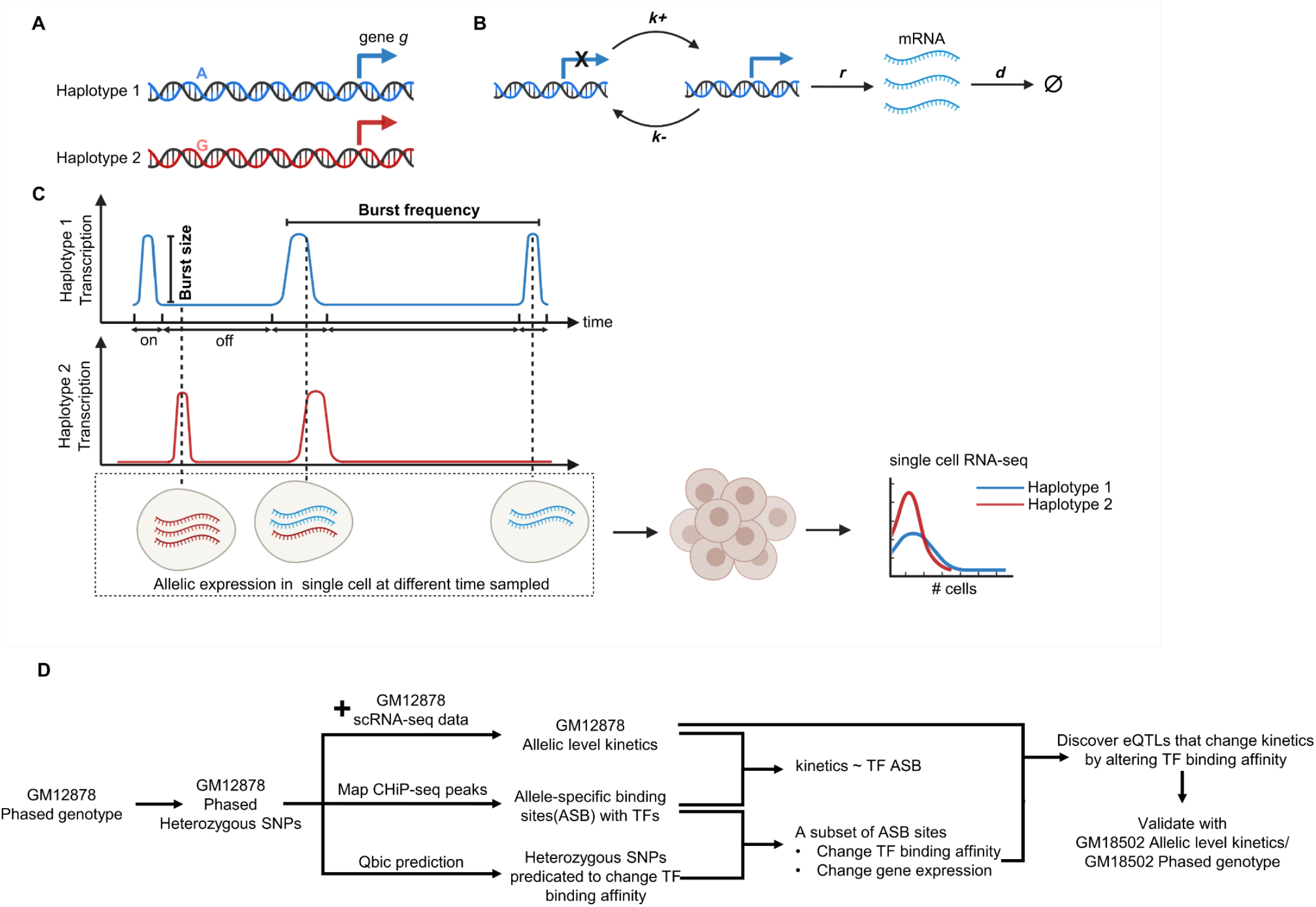
(A-C) Schema of the allelic transcriptional bursts. (A) Gene *g* has two promoters independently transcribing mRNA. (B) The promoter switches between an on and off state with rates k+ and k-. In the on state, a promoter can transcribe mRNA with rate *r*, which will be degraded with rate *d*. (C) The time-series transcription comprises episodic bursts of the transcript followed by silenced periods. The transcriptional bursts in single cells are the observed transcripts from Haplotype 1 or Haplotype 2. Different burst kinetics result in distinguished distributions of transcript between Haplotype 1 and Haplotype 2. (D) The overall analysis procedure for this manuscript.

**Figure 2.**
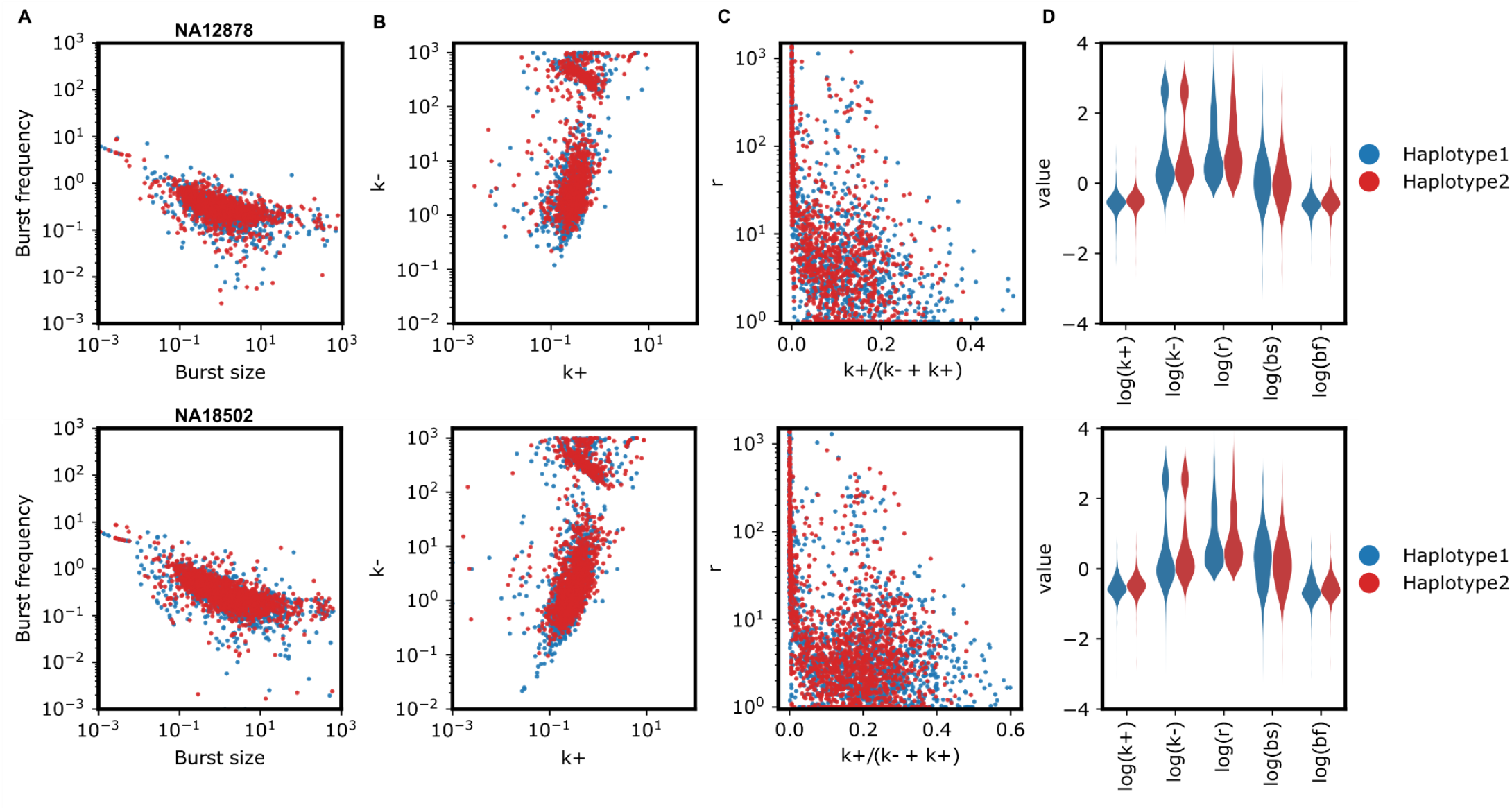
(A-D) The distribution of transcriptional kinetics for GM12878 and GM18502 (from up to down). Overall, there are 1,135 genes for GM12878 and 1,755 genes for GM5802. (A) shows the distribution of *k* and *k* in the log scale. (B) shows the distribution of *r* and 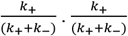 is related to the average active time fraction. (C) shows the distribution of burst size and burst frequency. The burst size and frequency are related to *k*_+_,*k*_−_, and *r* in the form of 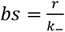 and 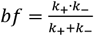. (D) The log-scaled violin plot of all transcriptional kinetics between Haplotype1 and Haplotype2.

### eQTL may exist that alter transcriptional kinetics rather than mean expression

Burst size and frequency can be independently regulated and lead to the change of allelic transcription. Typical eQTL studies investigate the association between genetic variants and mean expression in population. We wanted to determine if changes in mean expression is due to variants that alter transcriptional kinetics. If so, which burst kinetic in specific are altered by genetic variants.

We examined the correlation between our transcriptional kinetics and established sets of eQTLs from bulk expression experiments. We accessed 52,505 significant eQTL from LCLs across four studies in the eQTL Catalog (Kerimov et al., 2021). The effect sizes of these eQTLs are normalized to reflect the change of gene expression from Haplotype2 to Haplotype1. Then we correlated the eQTL effect size with the log-transformed fold change of transcriptional kinetics from Haplotype2 to Haplotype1 (see Methods).

We found that eQTLs affect gene expression mainly through altering k+ and burst frequency from the correlation results. First, the change of gene expression between alleles from the scRNA-seq data, quantified by mean and variance, establishes the highest correlation with the eQTL effect size (p-value=1.39e-42, Spearman correlation coefficient=0.336; p-value=9.57e-39, Spearman correlation coefficient=0.321). It is expected since eQTL effect sizes reflect the mean gene expression change between two alleles. Among transcriptional kinetics, the k+ and burst frequency correlate most with the eQTL effect size (p-value=4.57e-19, Spearman correlation coefficient=0.223; p-value=1.43e-15, Spearman correlation coefficient=0.2), respectively. Even though the eQTL sets are derived primarily from European descent subjects, we observe a similar correlation for k+ and burst frequency with eQTL effect size for GM18502 (Supplemental Figure S3).

Though k+ and burst frequency show the highest correlation with eQTL effect size, the correlation is only ∼0.2. Through a simulation study, we found that the missing correlation is likely because eQTL analysis cannot fully identify the change in gene expression when burst size and frequency are regulated. In a simulated case with 100 subjects and 1,000 cells per subject, we changed burst frequency and size simultaneously. In all simulated configurations, transcriptional kinetics analysis can identify the change in bursting kinetics, while eQTL analysis cannot (Figure 3B). Power estimates indicate that while transcriptional kinetics analysis can detect the change in burst kinetics with power greater than 60% with 18 subjects and 600 cells, eQTL analysis cannot detect expression changes. It is because when burst size and frequency are changed in opposite directions, the mean expression shows no statistical difference, while the variance does (Figure 3D). Therefore, when only a small number of cells are available, which is typical for tissue-based gene expression studies, it is more powerful to estimate transcription variance or transcriptional kinetics rather than mean expression.

**Figure 3.**
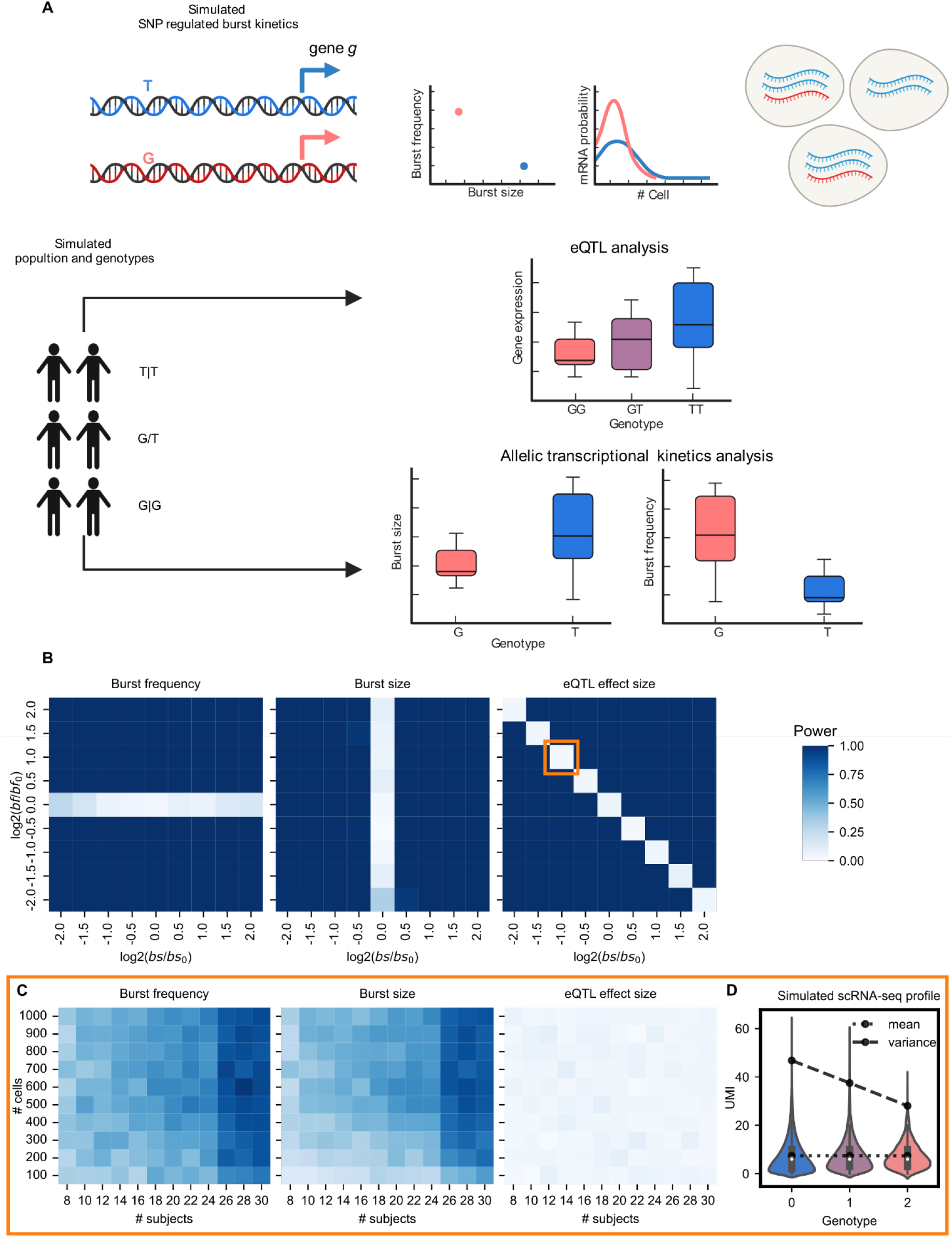
(A) The simulation workflow. A SNP will regulate gene *g* expression by modulating burst size and frequency. We simulated the genotype for n subjects and their allelic expression of gene g for M cells with Poisson-beta distribution with their burst size and frequency. The bulk gene expression for each subject is calculated by averaging through M cells. We then re-estimated the transcriptional kinetics based on each subject’s single-cell allelic expression. We compared the bursting kinetics for all subjects with the Student *t*-test. On the other side, we performed eQTL analysis on SNP J and bulk expression of gene g. (B) We simulated the experimentally ideal scenario, that is 100 subjects and 1,000 cells per subject, for eQTL and transcriptional kinetics analysis. The x-axis is the log_2_ transformed change from bs to *bs*_*0*_ and the y-axis is the log_2_ transformed change from bf to *bf*_*0*_ . Each combination of burst size and frequency was run 100 times independently. The power of either the eQTL test or the Student *t*-test of bursting kinetics is the percentage of tests reaching a significance level (alpha=0.05). (C) We simulated the scenario where burst size is *0*.*5bs*_*0*_ and burst frequency 2*bf*_*0*_, which is the scenario where eQTL analysis cannot detect the change in gene expression even with 100 subjects and 1,000 cells per subject. The x-axis is the number of samples from 8 to 30, and the y-axis is the number of cells from 100 to 1,000. Each combination of burst size and frequency was run 100 times independently. The power of either the eQTL test or the Student *t*-test of bursting kinetics is calculated by the percentage of tests reaching the significance level(alpha=0.05). (D) The violin plot of gene expression at single-cell resolution for different dosages of G alleles. The single-cell expression profile was simulated under the experimentally ideal scenario for both eQTL and transcriptional kinetics analysis with 100 subjects and 1,000 cells per subject. The amount of transcript from the T allele and G allele was simulated with bursting kinetics (*bs*_*0*_, *bf*_*0*_) and (*0*.*5bs*_*0*_,*2bf*_*0*_).

### Transcription factors bound to the transcription start site explain most transcriptional kinetics

We hypothesize that transcriptional kinetics are determined by features of the regulatory regions on the same chromosomal copy. To examine our hypothesis, we obtained the allelic level TF occupancy profile with ChIP-seq reads and phased genotype profiles and tested the association between allelic level TF occupancy and burst kinetics. It is worth noting that using complete phased genotype profiles is necessary and helpful to directly associate the TF binding event with genes on the same chromosomal copy.

First, we estimate the variance of transcriptional kinetics explained by the TF occupancy profile on the same phase with a linear mixed model (LMM). The TF occupancy profile is divided into the core promoter profile and the distal enhancer profile based on the regulatory regions that TFs bind (see Methods). In the core promoter profile, at most 90% of the variance of burst frequency can be explained by TF occupancy. However, only ∼30% variance for burst size is explained (Figure 4A). It is consistent with prior findings where burst frequency is highly regulated by transcription factors while burst size is not (Dobrinic et al., 2021; Fukaya et al., 2016). Compared to the core promoter profile, only 40% of the variance for burst size and 20% for burst frequency are explained by the distal enhancer profile. Therefore, transcriptional kinetics is largely contributed by TFs within the core promoter region.

**Figure 4.**
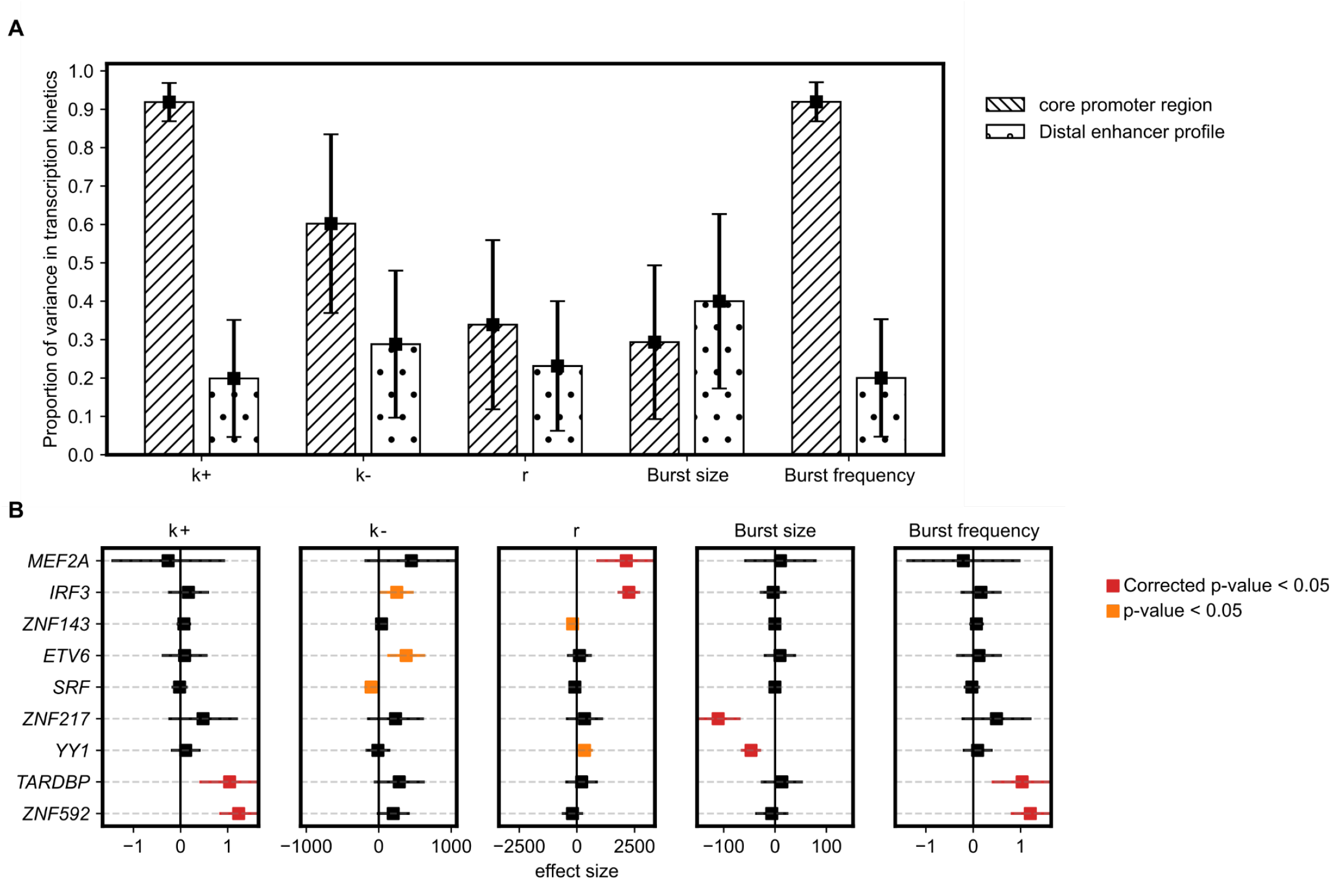
(A) The proportion of variance in transcriptional kinetics explained by TF binding. The bars with back slashes are the estimated variance of transcriptional kinetics explained by TF ASB within the core promoter, while the bars with dots are the estimated variance of transcriptional kinetics explained by TF ASB within the distal enhancer region. (B) Association results between the TF occupancy and transcriptional kinetics with a mixed multiple-variate linear model with gene as the group. Effect sizes that are significant with FDR<0.05 are colored red and those with nominal significance are colored orange.

Given that the TF ASB profile can explain a significant proportion of the variance in transcriptional kinetics on the same phase, we next want to identify the individual effect of TF on transcriptional kinetics. We combined the core promoter and distal enhancer ASB profiles and used a mixed linear model to investigate the individual TF effect in transcriptional kinetics. We found that the ASB events of multiple TFs are associated with transcriptional kinetics (Figure 4B). The binding of *ZNF217* and *YY1* in phase with the promoter is associated with gene expression by decreasing burst size. ASB of TFs binding in phase with the promoter, including *TARDBP* and *ZNF592*, are associated with gene expression by increasing burst frequency. *MEF2A* and *IRF3* increase the transcription rate *r* through TF ASB.

### eQTLs induce preferential TF binding that shapes the bursting kinetics of gene expression

In the eQTL section, we found that eQTLs affect average gene expression mainly through altering k+ and burst frequency from the correlation test. We also have shown that ASB of TF is associated with transcriptional kinetics. Here we combine them to infer potential eQTL mechanisms regulating burst kinetics by altering TF binding affinity and how they associate with phenotypes in the GWAS catalog.

The first two examples are eQTL loci. An eQTL rs4803042 of gene *ASB1* is carried by GM12878. The A allele of rs4803042 induces the preferential binding of *PORL2A* (Figure 5A), associating with a decreased burst size and increased burst frequency. rs4803042 overlaps with the *PORL2A* CHiP-seq peaks, which are pre-dominantly mapped to the rs4803042_A allele. In addition, the Qbic prediction (Martin et al., 2019) validates that the rs4803042_A allele has higher TF binding affinity than the rs4803042_G allele. In addition, we observe a similar increased burst frequency and decreased burst size for GM18502 at the rs4803042_A allele. Similar observations have been found in eQTL rs9266234 of gene *CCHCR1*. The T allele of rs9266234 induces the preferential binding of *POUF2F and TBP* (Figure 5B), associating with an increased burst size and decreased burst frequency. Even though rs9266234 has a flip genotype in GM18502, we observe a similar increased burst frequency and decreased burst size for GM18502 at the rs9266234_T allele.

**Figure 5.**
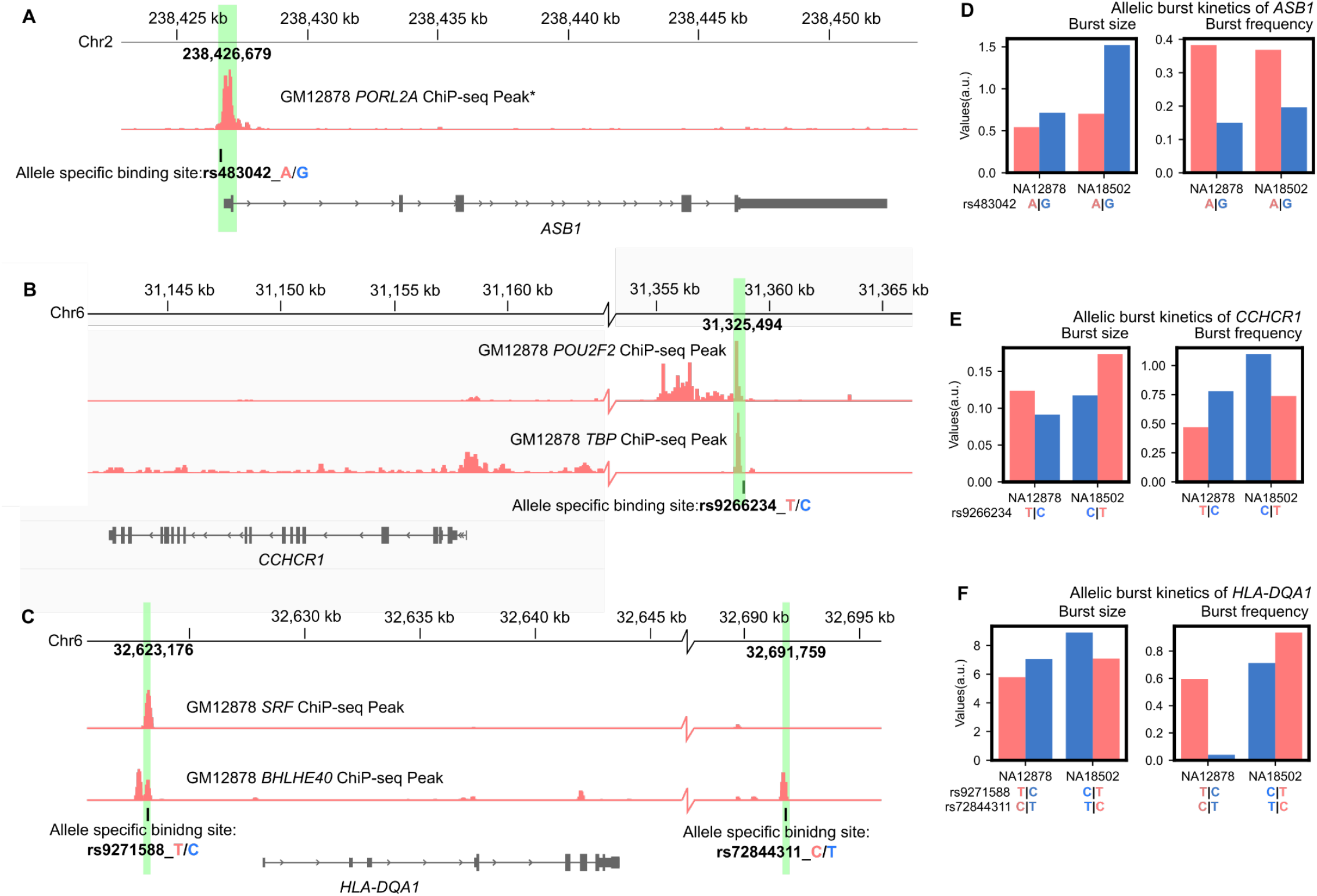
(A) rs4803042 is eQTL of gene *ASB1* carried by GM12878. The A allele of rs4803042 has shown ASB of *PORL2A* and is predicated to having higher TF binding affinity than the rs4803042_G allele. *The *PORL2A* ASB CHiP-seq results are replicated in two ENCODE experiments (ENCSR000DKT, ENCSR000BGD) (B) rs9266234 is eQTL of gene *CCHCR1* carried by GM12878. The A allele of rs9266234 has shown ASB of *POUF2F2*, and *TBP* is predicted to have higher TF binding affinity than the rs9266234_T allele. (C) rs9271588 and rs72844311 are eQTL of gene *HLA-DQA1* carried by GM12878. The T allele of rs9271588 has shown ASB of *SRF* and is predicated to having higher TF binding affinity than the rs9271588_C allele. The C allele of rs72844311, in phase with the rs9271588_T allele, also shows ASB of *BHLHE40* and is predicated to have higher TF binding affinity than its counter allele. (D-F) Comparison of the burst size and frequency between alleles and samples for gene *ASB1, CCHCR1*, and *HLA-DQA1*.

**Figure 6.**
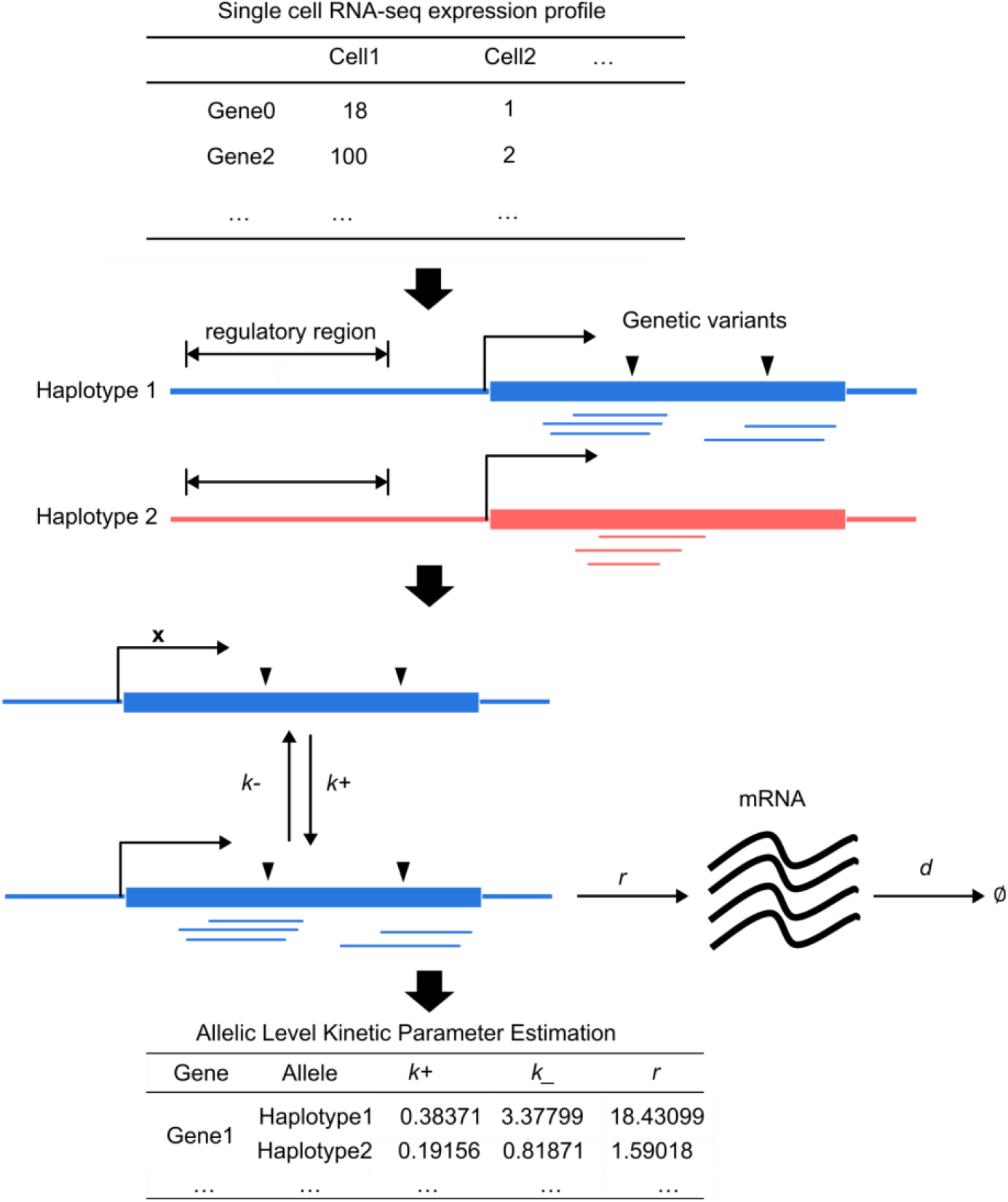
Diagram for kinetic parameter estimation from the scRNA-seq profile with phased genotype profile. The original scRNA-seq profile is quantified with unique molecular identifier (UMI). We mapped the scRNA-seq reads back to the chromosomes at heterozygous SNPs and derived the allelic level expression profile. Then the allelic level expression profile was fit with the two-state model for estimating three kinetic parameters: *k+, k-*, and *r*, which are scaled by the degradation rate *d*.

We observe an example rs9271588 that is both an eQTL and GWAS Catalog variant (Table 1a, b). The T allele of rs9271588 induces preferential binding of *SRF* and associate with higher burst frequency and larger transcription variance. rs9271588 is close to the transcription start site of *HLA-DQA1* and overlaps with the *SRF* ChIP-seq peaks, which are mostly mapped to the rs9271588_T allele (Figure 5C). The Qbic prediction confirms that the rs9271588_T allele has higher TF binding affinity. Moreover, another eQTL rs72844311 at downstream of *HLA-DQA1* was also shown to induce *BHLHE40* ASB at the C allele, which is in phase with the rs9271588_T allele. Again, we observe a similar increased burst frequency and decreased burst size for GM18502 at the haplotype rs9271588-rs9271588_TC. The rs9271588_C allele predicted to have lower TF binding affinity and, consequently, to result in low burst frequency is associated with an increased count of myeloid lineage cells. Lymphoid progenitor cells have similar HLA class II expression profiles to hematopoietic stem cells (Boegel et al., 2018). Therefore, rs9271588_C might have a similar influence on *HLA-DQA1* in hematopoietic stem cells to LCL, which results in cell-to-cell heterogeneity in hematopoietic stem cells and consequently an increased count of cells in the myeloid lineage.

**Table 1a:**
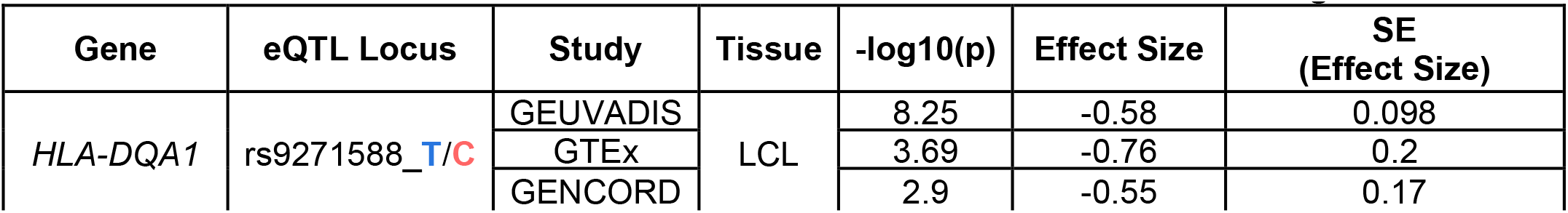
*HLA-DQA1* association with rs9271588 in the eQTL catalog

| Gene | eQTL Locus | Study | Tissue | -log10(p) | Effect Size | SE (Effect Size) |
| --- | --- | --- | --- | --- | --- | --- |
| *HLA-DQA1* | rs9271588_T/C | GEUVADIS | LCL | 8.25 | -0.58 | 0.098 |
|  |  | GTE<sub>x</sub> |  | 3.69 | -0.76 | 0.2 |
|  |  | GENCORD |  | 2.9 | -0.55 | 0.17 |

**Table 1b:** phenotypes for rs9271588,rs9271586 and rs9274623 in the GWAS catalog

| Variant and risk allele | P-value | Beta | CI | Trait(s) | Study accession |
| --- | --- | --- | --- | --- | --- |
| rs9271588-C | 2 x 10<sup>-58</sup> | 0.0576552 unit increase | [0.051-0.065] | neutrophil count, basophil count | GCST004620 |
|  | 2 x 10<sup>-66</sup> | 0.06164672 unit increase | [0.055-0.069] | granulocyte count | GCST004614 |
|  | 4 x 10<sup>-66</sup> | 0.06142442 unit increase | [0.054-0.068] | neutrophil count, eosinophil count | GCST004613 |
|  | 9 x 10<sup>-68</sup> | 0.06247162 unit increase | [0.055-0.07] | myeloid white cell count | GCST004626 |
|  | 1 x 10<sup>-58</sup> | 0.05764791 unit increase | [0.051-0.065] | neutrophil count | GCST004629 |

## Discussion

We systematically investigated the burst kinetics and their association with TF ASB in LCL. By associating the transcriptional kinetics with allelic-specific TF occupancy profile, we found that ASB of TF within the core promoter region explains 90% of burst size. Several individual TFs, including *YY1* and *SRF*, alter gene expression by changing burst size or frequency. Moreover, we identified multiple examples where eQTL or GWAS loci are ASB sites and are associated with the change in burst kinetics in GM12878. These examples are also replicated in GM18502. Our approach has the advantages that it resolves gene expression and burst kinetics at the allelic level and build the connection between ASE with coding variants and the regulatory effect from non-coding variant with haplotype information. In addition, our approach is a fine-mapping method to explain the molecular mechanism for eQTL results with a small sample size and a limited number of cells. In our simulation and real-life examples of eQTL loci, we demonstrate that there is a possibility that eQTLs exist by changing the burst kinetics but not the mean expression, which is challenging to be identified by typical eQTL studies.

We are aware that this study is limited to cell types, specifically lymphoblastoid cell lines. First, it is worthy emphasizing that GM12878 is the only few cell lines with more than 100 TF CHiP-seq, ATAC-seq, and DNA-methylation data available, which makes it the only few cell types to possibly investigate large-scale regulatory mechanisms in burst kinetics. Studies with similar ideas have been performed in K562 (Gupta et al., 2022) which, however, assumes a gene-level bursting mode and the large degree of aneuploidy within K562(Naumann et al., 2001; Zhou et al., 2019) and complicates allele-specific expression and perhaps promoter kinetic estimation.

Secondly, though here we have examined isogenic cell lines, we can apply our approach to typical scRNA-seq data by isolating homogenous cell types using multiple methods including marker gene expression, gene-gene co-concurrence (Qiu, 2020) and DNA methylation markers (Bianchi et al., 2022). To illustrate this idea, we tested our pipeline with a time-series scRNA-seq data on cardiac human pluripotent stem cell along differentiation (Friedman et al., 2018). By applying our method, we found multiple genes with significant change in transcriptional burst between pluripotent and differentiated cells (Supplemental Figure S4). These genes, unlike marker genes that completely turn on and off at pluripotent or differentiated states, are constantly expressed but with significantly different transcriptional variability at pluripotent or differentiated states due to the change in the burst kinetics. These genes are enriched in unique pathways for contractile and cardiac epithelial cells, further demonstrating the potential of our approach.

Two assumptions are made for the kinetic estimations: the ergodic process and constant mRNA degradation rate. The ergodic process assumption is valid for this study since no perturbation is introduced, and GM12878 is an isogenic cell line. The ergodic process can be violated when multiple cell types are involved, which can be resolved by characterizing the individual cell types. The second assumption is the constant mRNA degradation rate. Rabani et al. (2011) have shown that most genes in mammalian cells have a similar degradation rate, and the expression of those genes is little correlated with the change in degradation rates. Therefore, it is valid to estimate kinetics assuming a constant degradation rate and compare the transcriptional kinetics scaled by the degradation rate between genes.

The kinetic parameter estimation is sensitive to the protocol of scRNA-seq. Because the scRNA-seq dataset here is generated with the 10x Genomics 3’ sequencing, we have less coverage of exome regions with scRNA-seq reads and fewer allelic expression events identified. We use a stringent threshold to retain reads with unique alignment and high base-pair sequencing quality at the ASE SNP site. With such a strategy, we can estimate the allelic expression and transcriptional kinetics for 1,020 genes, while the original expression profile is available for 21,524 genes. We recognize that we compromise a larger proportion of lowly expressed genes during the QC process. However, more genes can likely be included in estimating the kinetic parameters with the latest scRNA-seq protocols, such as the Smart-seq3 (Hagemann-Jensen et al., 2020).

## MATERIALS AND METHODS

### Kinetic Parameter Estimation pipeline

#### 1) Obtain Allele-Specific expression from the scRNA-seq profile

The allele-specific mapping is done using the scRNA-seq data and phase genotype profiles. We first piled all scRNA-seq reads upon phased heterozygous SNPs with Samtools (Li et al., 2009). The minimum mapping quality for an alignment is 50, and the minimum base quality is 30. Reads mapped SNVs that are not in a single chromosome will be discarded. In detail, for GM12878, we found 7,375 cells with unique reads that have conflict SNVs, meaning that their SNVs are in different phases but appear in the same read. For GM18502, there are 6,429 cells with unique reads that have conflict SNVs.

Let *g* denote gene index, and *H1* denote Haplotype1. We processed cells with reads mapped to gene *g* and allelic-specific mapping information available. A read mapped to multiple SNPs on one chromosome within the gene *g* is counted as one single read. Reads mapped to multiple SNPs on both chromosomes are discarded. For convenience, these reads are termed reads with observed allelic origin. However, not all reads sequenced contain heterozygous SNVs. For these reads, we determine their allelic origin in a non-mapping way and term them as reads with inferred allelic origin.

Reads with observed allelic origin are used to estimate the ASE distribution with the following methods. Let 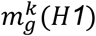 represents the number of transcripts from Haplotype1 for gene *g* in the *kth* cell and 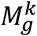 is the number of total transcripts for gene *g* in the *kth* cell. We assumed that in the *kth* cell with 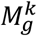 transcripts, the probability of observing 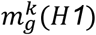 follows a beta-binomial distribution (1,2).

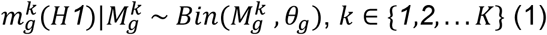

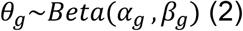

where *α*_*g*_ *and β*_*g*_ are canonical shape parameters of the beta distribution and *θ*_*g*_ is the probability sampled from the beta distribution. Based on *α*_*g*_ and *β*_*g*_, the expectation for observing a transcript from Haplotype1 within a single cell is shown in (3).

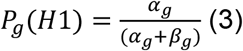

Let 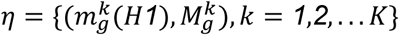 denotes the allelic expression observation in *K* cells. We first obtained the log-likelihood of the compound distribution shown in (4). We used the maximum likelihood estimation (MLE) to estimate the *α*_*g*_ and *β*_*g*_ on reads with observed allelic origin (5)

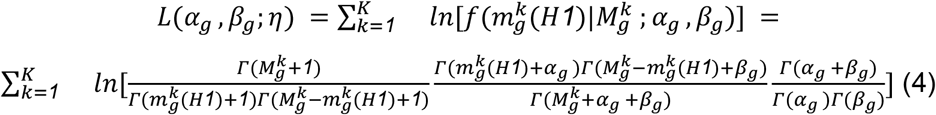

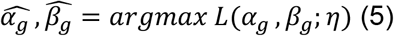

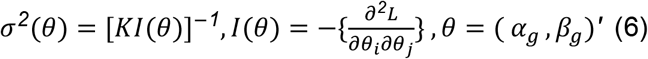

For cells with reads mapped to gene *g* but no allelic-specific reads available, we used the estimated beta-binomial distribution (1) and (2) infer the allelic transcript count of gene *g*. To ensure valid inference, we only keep genes with positive *α*_*g*_, *β*_*g*_, and confidence intervals. *α*_*g*_ and *β*_*g*_ estimated with an unrealistic confidence interval (i.e., confidence interval covering 0) indicates a failed convergence during maximum likelihood estimation and thus, will not be kept.

In summary, with either observed allele-specific reads or inferred allele-specific reads, we divided the overall unique molecular identifier (UMI) to Haplotype1 and Haplotype2 and consequently obtained the allele-specific expression profile for gene *g*.

#### 2) Estimate Kinetic Parameters

The stochastic gene expression is described by a two-state model (Figure 8). In the two-state model, a gene switches between active and inactive status with rates of *k+, k-*. While in the active status, the gene can transcribe mRNA with a rate of r. The degradation rate of a transcript is *d*.

**Figure 7.**
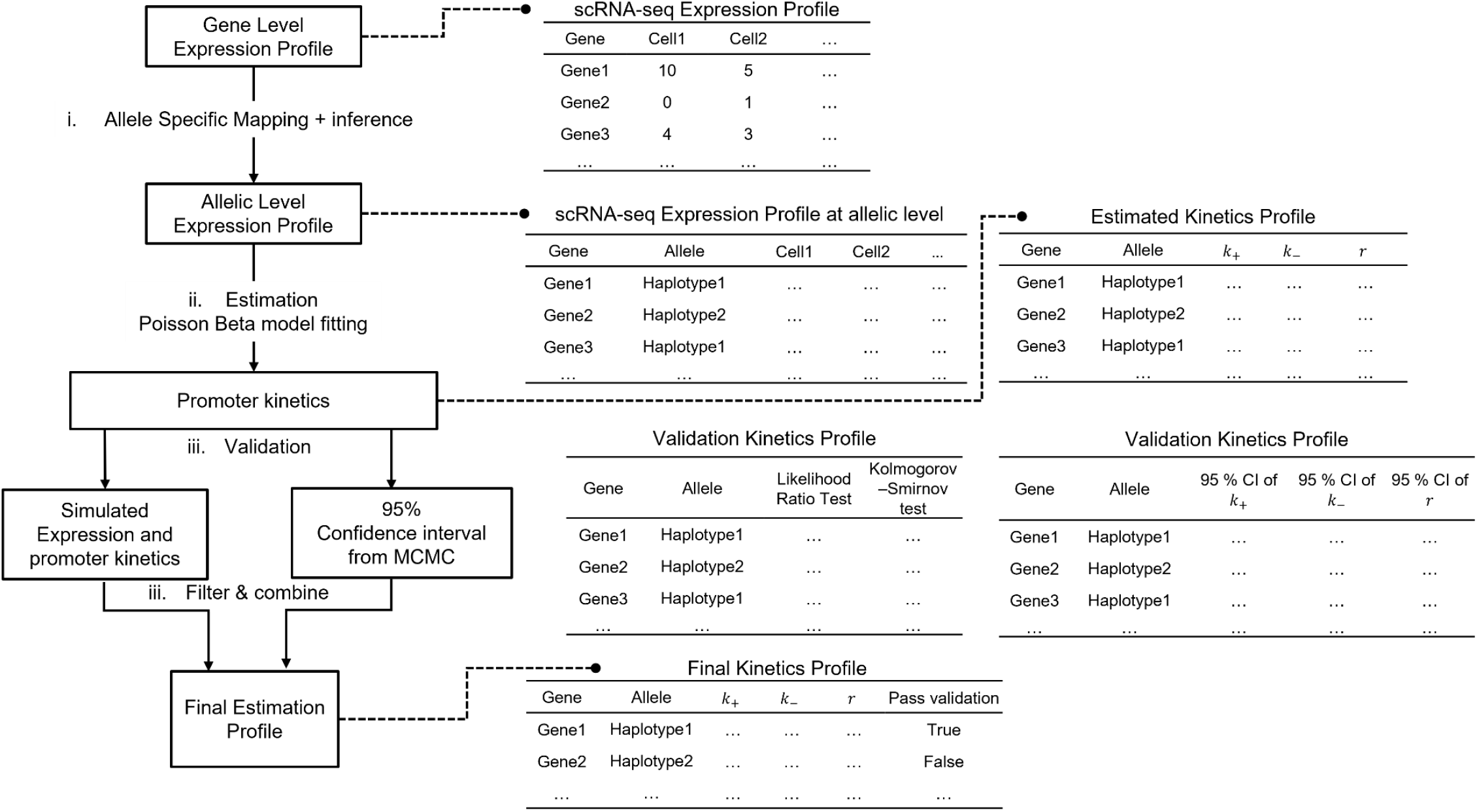
Workflow for the allelic-specific mapping, kinetic estimation, and validation process.

**Figure 8.**
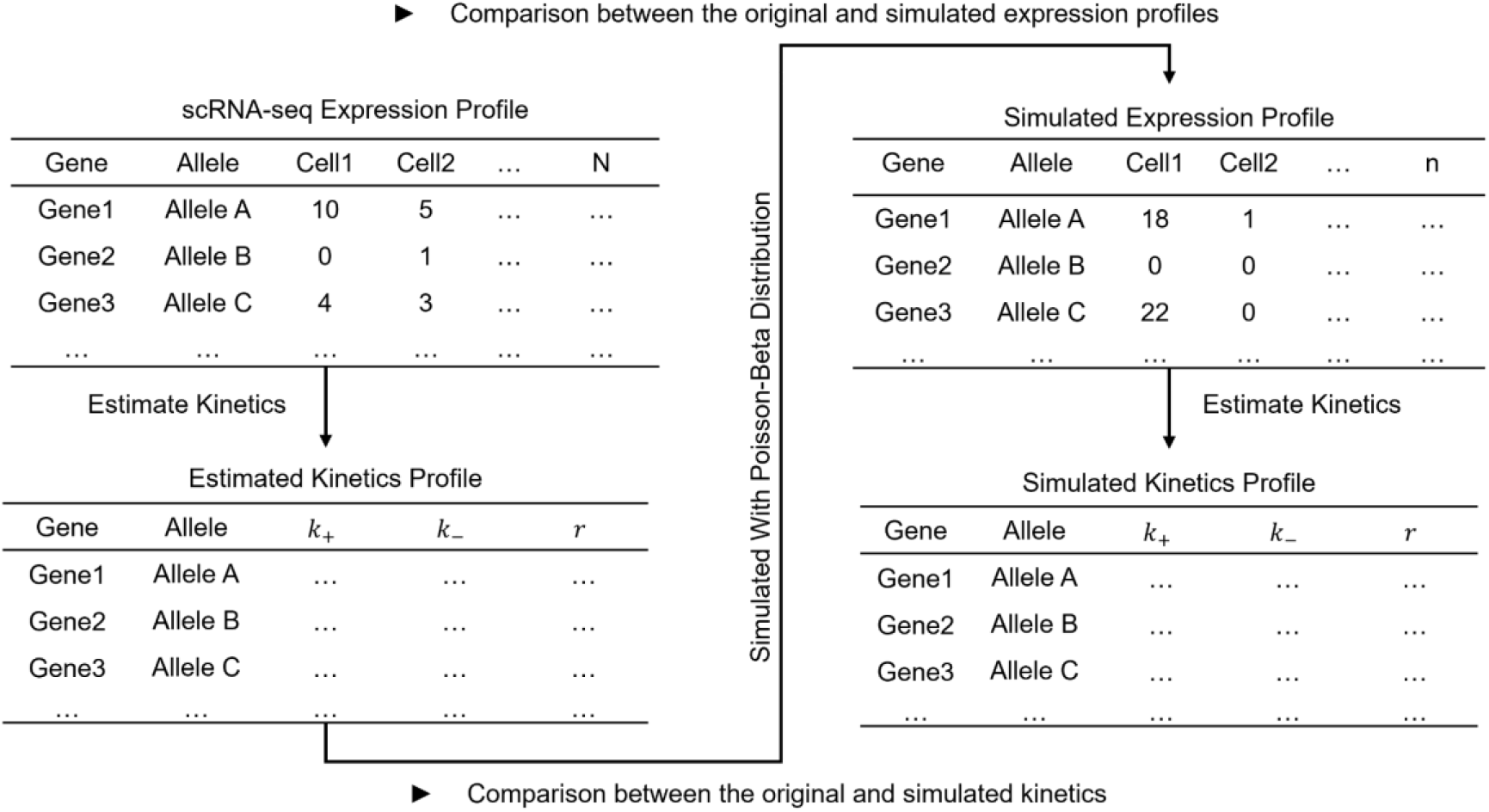
The flow chart of the first validation approach

We used the Poisson-Beta distribution as the steady-state distribution for the two-state model and estimated the kinetic parameters with the maximum likelihood approach. The probability density function for Poisson-beta distribution takes the same form as the analytical solution for the two-state model (Bressloff, 2014). We denote *m* as the number of mRNA in the cell, *m*_*s*_ = *r*/*d* is the mRNA number at the steady-state, *ξ*_*0*_ = *k*_−_/*d, ξ*_*1*_ = *k*_+_/*d*, and *F* is the confluent hypergeometric function of the first kind (Shahrezaei & Swain, 2008) The analytical solution for the two-state model is in Formula 7. The Poisson-beta distribution for a representative steady-state distribution is shown in Formula 8 (Vu et al. 2016), where *λ* is the hyper-parameter can be interpreted as the stochastic switch rate.

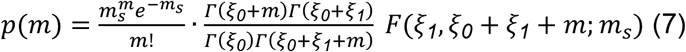

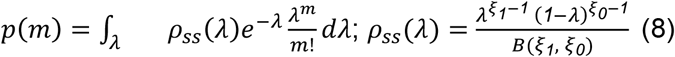

The kinetic parameters estimated from are *k+, k-*, and *r*, scaled by *d*. However, *k+, k-*, and *r* are not necessarily independent. Therefore, we derived a set of empirically orthogonal bursting kinetics, including bursting size (bs) and burst frequency (bf).

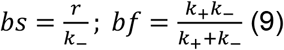

#### 3) Validate, Filter, and Generate Valid Kinetic Parameter Profile

We validated the estimated kinetic parameter profile with two independent approaches and kept estimated kinetic parameters whose validation profile has passed either filtering criteria.

In the first approach, we simulated gene expression based on the kinetic parameters estimated from the original expression profile. We then re-estimated the kinetic parameters again from the simulated expression profile. This approach examines both the correlation between the original and simulated expression profiles and the correlation between the estimated and simulated kinetic parameter profiles. We used the Kolmogorov–Smirnov test and the likelihood ratio test to test the correlation between the original and simulated expression profiles. Kolmogorov–Smirnov and Likelihood ratio tests being insignificant indicates that the underlying probability distributions for the original expression profile do not deviate from the simulated expression profile. We also require the estimated kinetic parameter 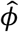 and the simulated kinetic parameter *ϕ*_*s*_to be restricted by 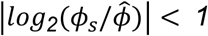, indicating that the simulated kinetic parameters do not differ from the originally estimated kinetic parameters. In summary, estimated kinetic parameters are kept if Kolmogorov–Smirnov and Likelihood ratio tests are not significant and the estimated kinetic parameter 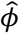 and the simulated kinetic parameter are restricted by 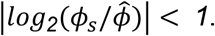.

In the second approach, we measure the variance of each estimated kinetic parameter and filter out those with large variance. Let *M* = {*m*_*k*_, *k* = *1,2*, . . . *K}* denote the observation of transcript in K cells. We sampled the posterior distribution *p*(*k*_+_, *k*_−_, *r*|*M*) with the Metropolis-Hastings algorithm (Zoller et al., 2018) in 10,000 iterations. The 95% credible interval are derived from the marginal posterior distribution. We require the upper bound 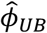 and lower bound 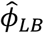 of the confidence intervals for estimated kinetic parameters to be restricted by 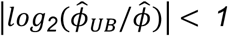 and 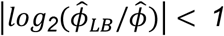.

### Allelic binding of transcriptional factors and histone modification

We combine the ChIP-seq data and phased genotype from GM12878 to identify allelic-specific TF binding and histone modification events, generalized as ASB in the following. We used 142 TF and 8 histone ChIP-seq datasets for GM12878 from the ENCODE4 project (Davis et al., 2018). We realigned the ChIP-seq reads upon heterozygous SNPs located in the regulatory region and tested if there were allelic-specific binding events with BaalChIP (de Santiage et al., 2017). The regulatory regions are defined as the distal enhancer region from EnhancerAtlas (Gao & Qian, 2020) and the core promoter region with 1,000 base pairs (bp) upstream and downstream from the transcription start site. The distance between the boundary of the enhancer region and the gene ranges from 1,001bp to 9.7Mbp, with a mean of 1.4M. SNPs are clustered if they are located within the same CHiP-seq peak. If within a single CHiP-seq peak, there are multiple SNPs showing both ASB and nonASB events, this CHiP-seq peak will not be considered to have an ASB event.

We derived the TF occupancy profile from ChIP-seq data by encoding the ASB events. Based on the phased genotype and TF allelic reads mapping for SNPs that show ASB, we determined if the TF preferably binds to Haplotype1 or Haplotype2. The preferably bound allele is encoded as 1 for TF occupancy, while the counterpart allele is encoded as 0. If a single CHiP-seq peak shows conflict ASB events by multiple SNPs, it will not be considered an ASB event.

### Associate the transcriptional kinetics with the eQTL effect size

We accessed 935,536 significant eQTL from LCLs across four studies in the eQTL Catalog (Kerimov et al., 2021). We retained 52,505 eQTL variants that were heterozygous in GM12878 and were potential eQTLs (alpha=1e-05) for genes with estimated kinetics.

Given that the eQTL effect size defaults as the change from the alternative allele against the reference allele, we use the following strategy to make the change of transcriptional kinetics comparable with the change of eQTL between alleles. First, based on the phased genotype, we normalized the eQTLs effect size to reflect the change of gene expression from Haplotype2 to Haplotype1. Meanwhile, we derived the log-transformed fold change of transcription burst from Haplotype2 against Haplotype1. With allelic transcriptional kinetics change comparable to the change of eQTL effect size between alleles, we tested the association between them with the spearman correlation test.

### Power simulation

In Figure 3B, we simulate the scenario for both eQTL and transcriptional kinetics analysis with *N*=100 subjects and *K*=1,000 cells per subject. We choose the bursting kinetics *bs*_*0*_ and *bf*_*0*_ from *HLA-DQA1* estimated from GM12878. We simulate SNP *J*, the allele of which results in two different sets of bursting kinetics (*bs, bf*) and (*bs*_*0*_, *bf*_*0*_) for gene *g*.

We determined the minor allele frequency *p* for the regulatory SNP *J* and simulated the genotype for *N*=100 subjects assuming SNP *J* following Hardy-Weinberg equilibrium. Based on their genotype, we simulated each subject expression of gene *g* for *K=*1,000 cells. 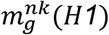 is the expression of Haplotype1 allele of gene *g* for the *n*th subject in the *k*th cell, which follows a Poisson-beta distribution with their burst size 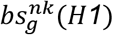 and burst frequency 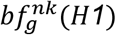 (10). Similar annotation applies to 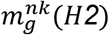, 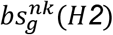, and 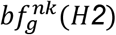. The bulk expression *g*_*n*_ for the *n*th subject is obtained by averaging across *K* cells and two alleles H1 and H2 (12). We then re-estimated the transcriptional kinetics for the *k*th subject based on its allelic expression profile 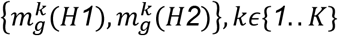 across K cells (13, 14). We compared the bursting kinetics for all subjects with the Student *t*-test. On the other side, we perform eQTL analysis on SNP *J* and the bulk expression of gene *g*. Finally, we compared the results from the eQTL analysis and the re-estimated bursting kinetics. Each combination of burst size and burst frequency was run 100 times independently. The power of either the eQTL or burst kinetic test is the percentage of tests reaching the significance level (alpha=0.05).

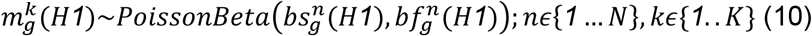

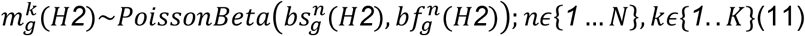

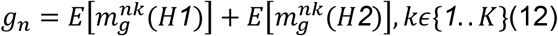

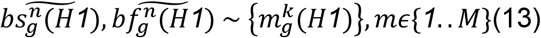

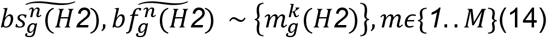

Similarly for Figure 4C, we set up the scenario where the allele of SNP *J* will result in a corresponding set of bursting kinetics (*0*.*5bs*_*0*_, *2bf*_*0*_) and (*bs*_*0*_ and *bf*_*0*_). We varied the number of samples *N* from 8 to 30 and the number of cells *K* from 100 to 1,000. Each combination of *K* and *N* was run 100 times independently. The power of either the eQTL or burst kinetic test is calculated by the percentage of tests reaching the significance level (alpha=0.05).

### Associate the transcriptional kinetics with the ASB profile

We used a linear mixed model to estimate the variance of transcriptional kinetics explained by TF occupancy. Let *D* denotes the TF occupancy profile and *σ*^*2*^ as the variance of a Transcriptional kinetic explained by *D*.

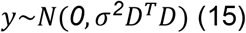

While LMM requires the residual to follow a normal distribution, we did not know if the transcriptional kinetics violate the assumption. However, we argue that we only use the LMM to estimate the strength of the association between transcriptional kinetics and TF occupancy. Suppose the TF occupancy can explain a large proportion of the variance of transcriptional kinetics. In that case, we have theoretical evidence to support transcriptional kinetics are likely contributed by gene regulation, which allows us to further investigate the individual TF and their effect on transcriptional kinetics.

We combined the core promoter and distal enhancer ASB profiles to investigate the individual TF effect in transcriptional kinetics. Since both core promoter and distal enhancer regions have ASB profiles combining them helps increase the number of ASB events and, thus, the statistical power. There are 607 (607/1,135) genes with ASB events within either the core promoter or distal enhancer region, and these ASB events come from 107 (107/150) TFs and Histone markers. We performed a mixed multiple variate linear model with gene as the group. A positive test effect size indicates that the existence of a TF or Histone will increase the transcriptional kinetic and vice versa. We correct the test result for multiple testing with the False Discovery Rate (FDR).

### scRNA-seq data source and quality control (QC)

We used two criteria to remove abnormal cells and maintain cell-cell variability for the scRNA-seq expression profile. Firstly, we preserved cells with <10% of the transcript from mitochondrial genes. In addition, cells with fewer UMI than 2.5 standard deviations from the population means are excluded.

The scRNA-seq data for GM12878 and GM18502 is from Osorio et al. (2019). For GM12878, 8,287 cells are sequenced, and 7,817 pass the quality control (QC). Using these 7,817 cells from GM12878, we obtain the valid estimation of transcriptional kinetics for 2,184 alleles corresponding to 1,135 genes. For GM18502, 5,991 cells are sequenced, and 5,749 pass the QC. We obtain the valid estimation of transcriptional kinetics for 1,755 genes with 3,377 alleles.

## SOFTWARE AVAILABILITY

The code for this study is available on GitHub: https://github.com/bushlab-genomics/ASEkinetics with open access or can be found in the Supplemental code file.

## ACKNOWLEDGMENTS

This work is supported by U01 AG058654 (Haines, Bush, Martin, Farrer, Pericak-Vance) and RF1 AG070935 (Griswold, Bush)

Figures are created with BioRender.com.

## Supplementary Material

Despite the different continental ancestry of the two individuals, the single-cell allelic expression for these two LCLs is highly consistent. For GM12878, 1,008 out of 1,135 genes with an average probability of having a transcript from Haplotype1 around 0.5 are from a bimodal distribution (Figure S1C). Similarly, for GM18502, the majority of the genes (1,566 out of 1,755) expressed with bimodal distributions (Figure S1F). Genes with bimodal distributions are expressed preferably with one of the two alleles within a single cell, which is different from genes with unimodal distribution expressing both alleles within a single cell, even though both distributions can result in biallelic expression in the population average.

**Supplemental Figure S1.**
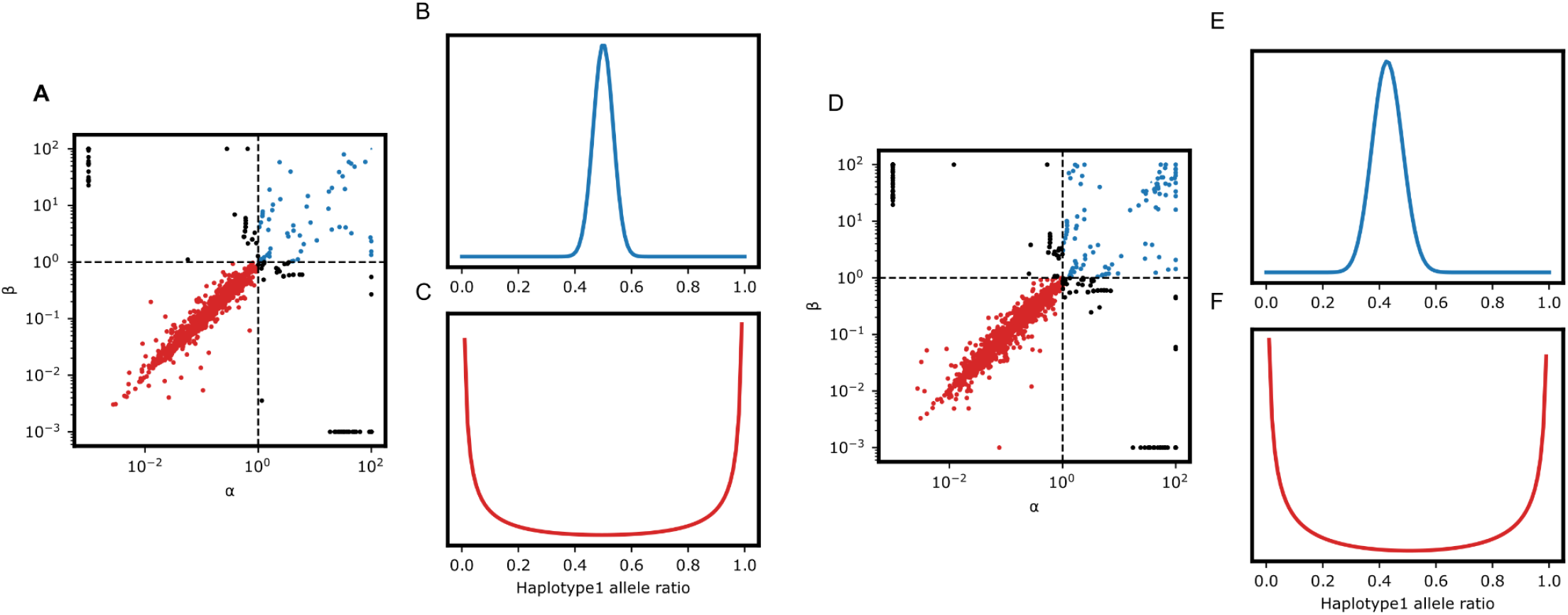
The allelic expression pattern for GM12878 and GM18502. (A) Distribution of α and β for GM12878. α and β are parameters that control the shape of the beta-binomial distribution. (B) The theoretical curve with α and β larger than 1 (C) The theoretical curve based on the estimated α and β smaller than 1. B and C show direct compassion between bimodal and unimodal distributions, both of which can result in biallelic expression autosomes. (D) Distribution of estimated α and β for GM18502. (E) The theoretical curve with estimated α and β larger than 1. (F) The theoretical curve with α and β smaller than 1.

**Supplemental Figure S2.**
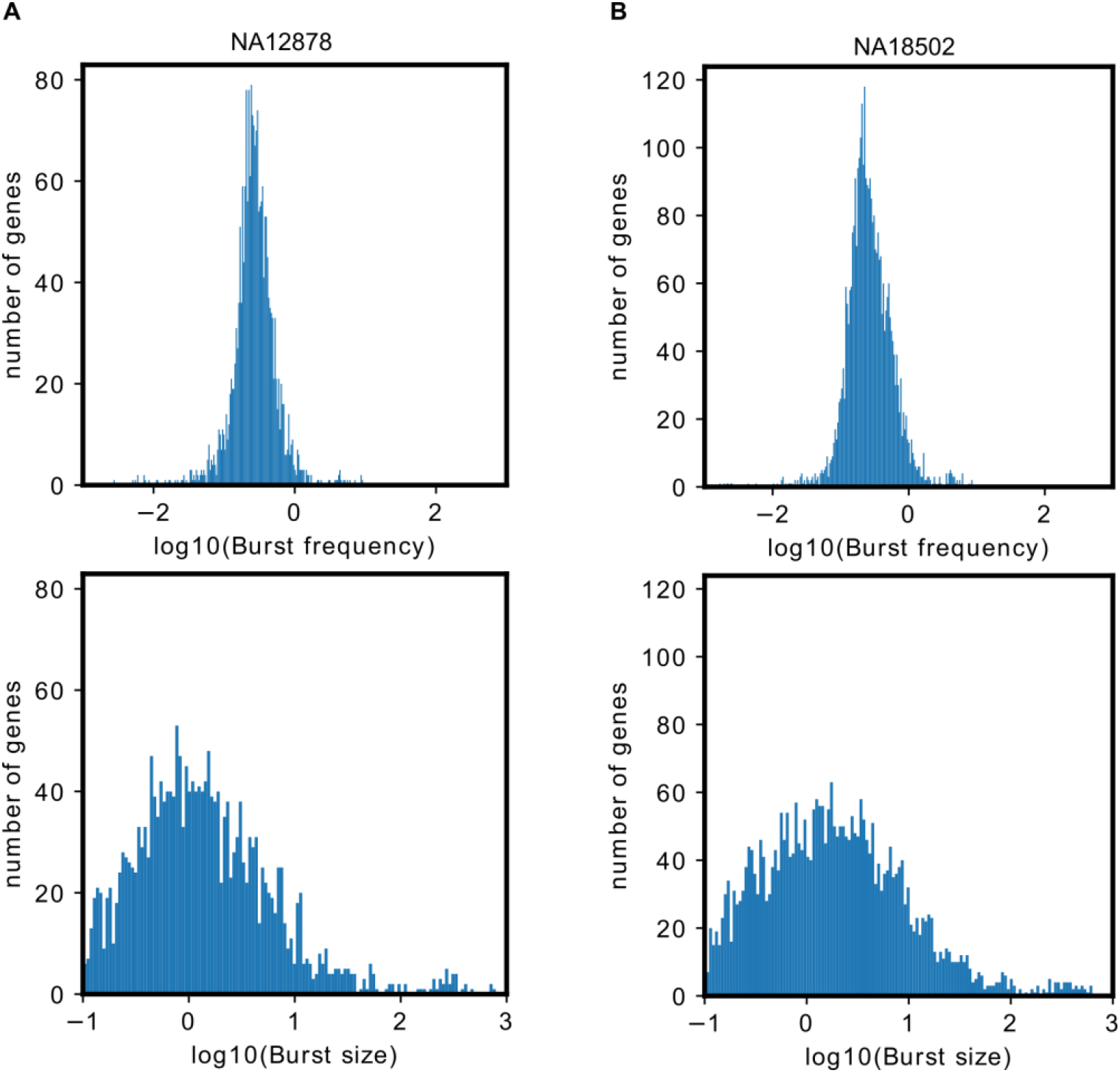
The histogram of log-transformed bursting kinetics for GM12878 and GM18502

**Supplemental Figure S3.**
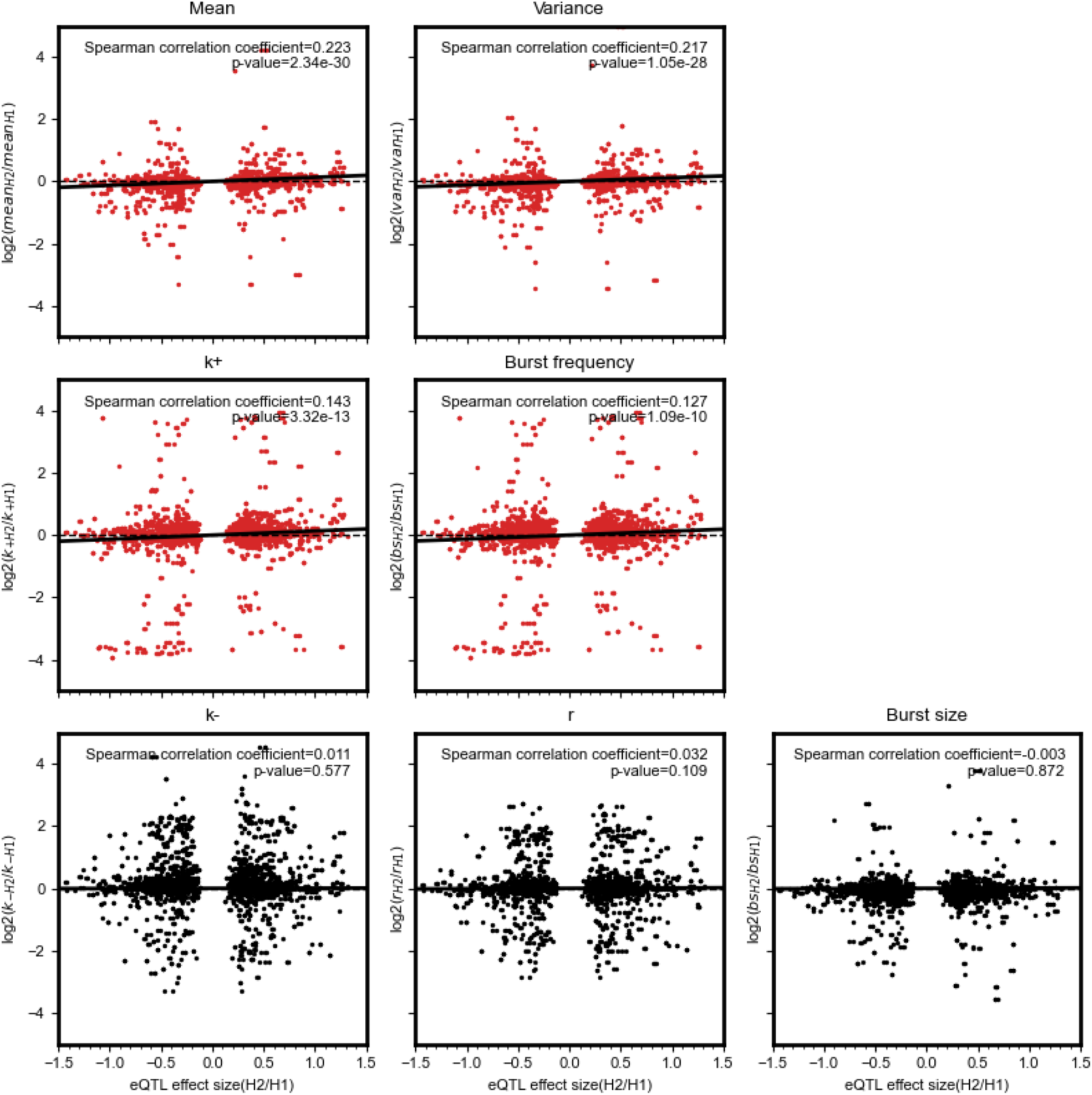
(A-G) Spearman correlation between eQTL effect size and transcriptional kinetics for GM18502. The x-axis is the effect size from the previous eQTL studies, and the y-axis is the log_2_ fold change of transcriptional kinetics. The directions for both eQTL effect size and log fold change of transcriptional kinetics are from Haplotype2 to Haplotype1. (A-B) The mean and variance change between alleles have the highest correlation with the eQTL effect size. (C-D) k+ and burst frequency exhibit the largest correlation coefficient with the eQTL effect size. (E-G) k-, r, and burst size are not correlated with the eQTL effect size.

### Applying the kinetic parameter estimation pipeline on other cell types

To demonstrate that our approach applies to general single-cell RNA-seq studies with multiple cell types, we applied it to an independent dataset from Friedman et al. (2018). This dataset is designed to study changes in transcriptomic profiles along cardiac differentiation of human pluripotent stem cells. We specifically examined day 0 when all cells maintain pluripotency and day 30 when cells are at definitive cardiac cell states. On day 0, the pluripotency gene *POU5F1* was expressed in the entire cell population and there are no distinguished cell types (Supplemental Figure S4A). On day 30, *MYL7* expression clearly distinguishes subpopulation day30:S1 from subpopulation day30:S2 (Supplemental Figure S4B).

We estimate the transcriptional kinetics for day 0 as a homogeneous cell group and separately for day30:S1 and day30:S2 for day 30. We compare genes that are different in burst kinetics in S1 and S2 against day 0. This analysis is designed to discover genes that change their expression kinetics along the differentiation. These genes, unlike *POU5F1* and *MYL7* which are either expressed or not on day 0 and day 30, were constantly expressed but with different kinetics and expression variability on day 0 and day 30. The change in the transcriptional variability has been implicated in pluripotency (Chang et al. 2008; Kalmar et al. 2009; Ohnishi et al. 2014).

We found unique pathways with enrichment of genes with significant changes in burst kinetics in day30:S1 and day30:S2 compared to day 0. Genes enriched in actin binding and cytoskeletal protein binding have significant fold change in burst kinetics in day30:S2 as contractile cells. Known cardiac sarcomere genes including *MYH6* and *MYH7B* have around 4 times change in the burst size, indicating their higher expression variability in day30:S1 compared to Day 0.

On the other side, genes enriched in cell mobility and locomotion have at least two fold change in burst kinetics in day30:S1 cells, which have similar expression profile to cardiac outflow tract (OFT) (Friedman et al., 2018). Cardiac endothelial cells are part of the OFT and their expression profile is found enriched in multiple cell migration related pathways (Lother et al. 2018), which correlates well with our results. Therefore we conclude that the change in burst kinetics and variability is associated with the cardiac cell state.

**Supplemental Figure S4.**
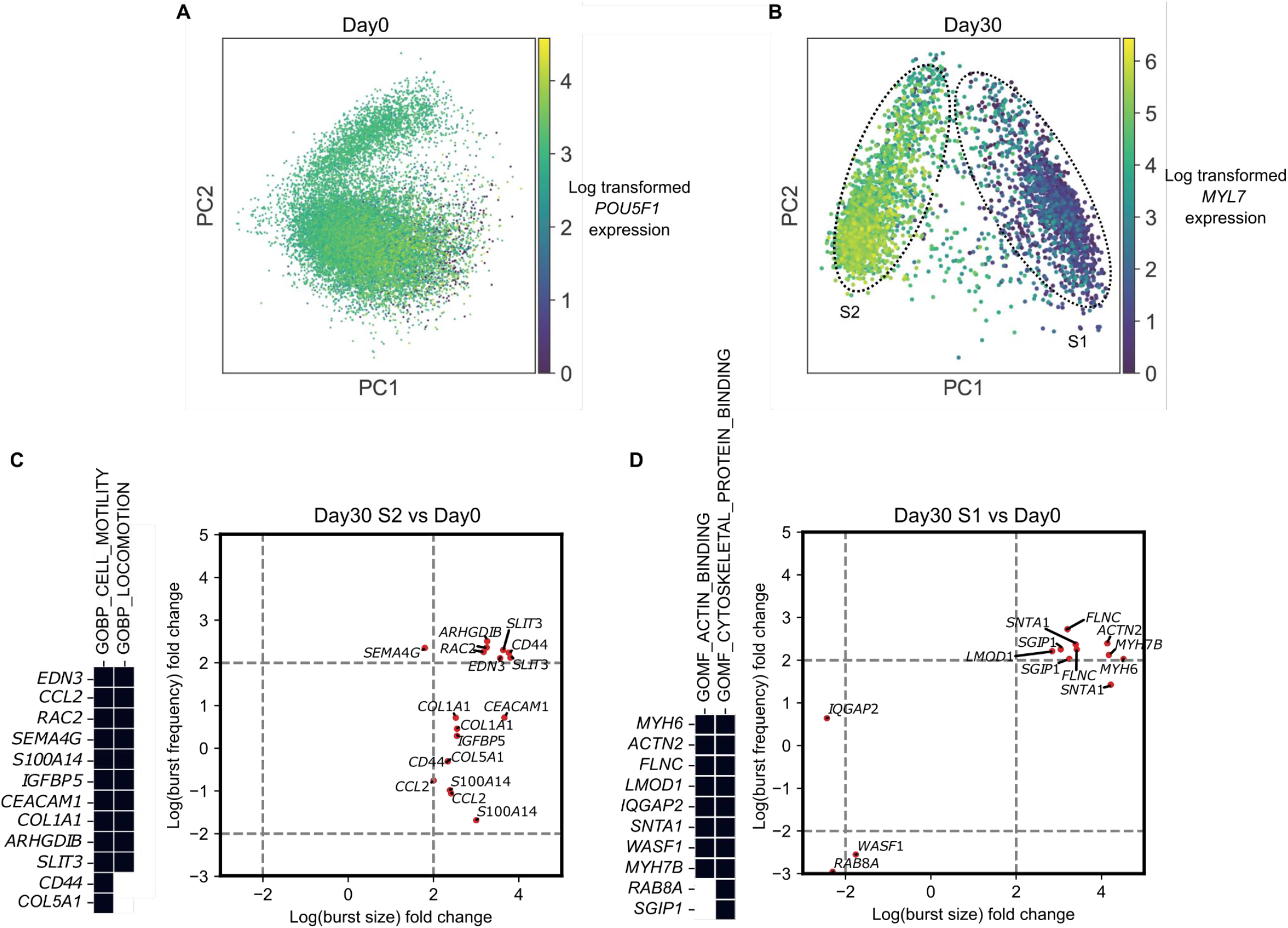
Transcriptional kinetics along cardiac differentiation of human pluripotent stem cells. A. Principle component analysis of human pluripotent stem cells at day 0. *POU5F1* is pluripotency gene expressed across the cells. B. Principle component analysis of human pluripotent stem cells at day 30. *MYL7* expressed differentially between subpopulation S1 and subpopulation S2. C. Genes enriched in cell mobility and locomotion pathway have more than 2-fold change in the burst kinetics when comparing day30:S2 against day 0. D. Genes enriched in actin binding and cytoskeletal protein binding have more than 2-fold change in the burst kinetics when comparing day30:S1 against day 0.

## REFERENCE

Kærn M, Elston TC, Blake WJ, Collins JJ. 2005. Stochasticity in gene expression: from theories to phenotypes. Nat Rev Genet 6: 451–464.

Bartman CR, Hamagami N, Keller CA, Giardine B, Hardison RC, Blobel GA, Raj A. 2019. Transcriptional Burst Initiation and Polymerase Pause Release Are Key Control Points of Transcriptional Regulation. Molecular Cell 73: 519-532.e4.

Prashant NM, Alomran N, Chen Y, Liu H, Bousounis P, Movassagh M, Edwards N, Horvath A. 2021. SCReadCounts: estimation of cell-level SNVs expression from scRNA-seq data. BMC Genomics 22: 689.

Borel C, Ferreira PG, Santoni F, Delaneau O, Fort A, Popadin KY, Garieri M, Falconnet E, Ribaux P, Guipponi M, et al. 2015. Biased Allelic Expression in Human Primary Fibroblast Single Cells. The American Journal of Human Genetics 96: 70–80.

Kumar RM, Cahan P, Shalek AK, Satija R, Jay DaleyKeyser A, Li H, Zhang J, Pardee K, Gennert D, Trombetta JJ, et al. 2014. Deconstructing transcriptional heterogeneity in pluripotent stem cells. Nature 516: 56–61.

Chang HH, Hemberg M, Barahona M, Ingber DE, Huang S. 2008. Transcriptome-wide noise controls lineage choice in mammalian progenitor cells. Nature 453: 544–547.

Gulati GS, Sikandar SS, Wesche DJ, Manjunath A, Bharadwaj A, Berger MJ, Ilagan F, Kuo AH, Hsieh RW, Cai S, et al. 2020. Single-cell transcriptional diversity is a hallmark of developmental potential. Science 367: 405– 411.

Jiang Y, Zhang NR, Li M. 2017. SCALE: modeling allele-specific gene expression by single-cell RNA sequencing. Genome Biology 18: 74.

Kim JK, Marioni JC. 2013. Inferring the kinetics of stochastic gene expression from single-cell RNA-sequencing data. Genome Biology 14: R7.

Bahar R, Hartmann CH, Rodriguez KA, Denny AD, Busuttil RA, Dollé MET, Calder RB, Chisholm GB, Pollock BH, Klein CA, et al. 2006. Increased cell-to-cell variation in gene expression in ageing mouse heart. Nature 441: 1011–1014.

Lammers, N.C., Kim, Y.J., Zhao, J., and Garcia, H.G. (2020). A matter of time: Using dynamics and theory to uncover mechanisms of transcriptional bursting. Current Opinion in Cell Biology 67, 147–157.

Ohnishi Y, Huber W, Tsumura A, Kang M, Xenopoulos P, Kurimoto K, Oles AK, Araúzo-Bravo MJ, Saitou M, Hadjantonakis A-K, et al. 2014. Cell-to-cell expression variability followed by signal reinforcement progressively segregates early mouse lineages. Nat Cell Biol 16: 27–37.

Martinez-Jimenez CP, Eling N, Chen H-C, Vallejos CA, Kolodziejczyk AA, Connor F, Stojic L, Rayner TF, Stubbington MJT, Teichmann SA, et al. 2017. Aging increases cell-to-cell transcriptional variability upon immune stimulation. Science 355: 1433–1436.

Rodriguez J, Larson DR. 2020. Transcription in Living Cells: Molecular Mechanisms of Bursting. Annual Review of Biochemistry 89: 189–212.

Dar RD, Razooky BS, Singh A, Trimeloni TV, McCollum JM, Cox CD, Simpson ML, Weinberger LS. 2012. Transcriptional burst frequency and burst size are equally modulated across the human genome. Proceedings of the National Academy of Sciences 109: 17454–17459.

Sánchez Á, Kondev J. 2008. Transcriptional control of noise in gene expression. PNAS 105: 5081–5086.

Nicolas D, Zoller B, Suter DM, Naef F. 2018. Modulation of transcriptional burst frequency by histone acetylation.

Proc Natl Bartman CR, Hamagami N, Keller CA, Giardine B, Hardison RC, Blobel GA, Raj A. 2019. Transcriptional Burst Initiation and Polymerase Pause Release Are Key Control Points of Transcriptional Regulation. Molecular Cell 73: 519-532.e4. Acad Sci USA 115: 7153–7158.

Antolović V, Miermont A, Corrigan AM, Chubb JR. 2017. Generation of Single-Cell Transcript Variability by Repression. Current Biology 27: 1811-1817.e3.

Li C, Cesbron F, Oehler M, Brunner M, Höfer T. 2018. Frequency Modulation of Transcriptional Bursting Enables Sensitive and Rapid Gene Regulation. Cell Systems 6: 409-423.e11.

Faure AJ, Schmiedel JM, Lehner B. 2017. Systematic Analysis of the Determinants of Gene Expression Noise in Embryonic Stem Cells. Cell Systems 5: 471-484.e4.

Fukaya T, Lim B, Levine M. 2016. Enhancer Control of Transcriptional Bursting. Cell 166: 358–368.

Singh A, Razooky B, Cox CD, Simpson ML, Weinberger LS. 2010. Transcriptional Bursting from the HIV-1 Promoter Is a Significant Source of Stochastic Noise in HIV-1 Gene Expression. Biophysical Journal 98: L32–L34.

Dobrinić P, Szczurek AT, Klose RJ. 2021. PRC1 drives Polycomb-mediated gene repression by controlling transcription initiation and burst frequency. Nat Struct Mol Biol 28: 811–824.

Larsson AJM, Johnsson P, Hagemann-Jensen M, Hartmanis L, Faridani OR, Reinius B, Segerstolpe Å, Rivera CM, Ren B, Sandberg R. 2019. Genomic encoding of transcriptional bursting kinetics. Nature 565: 251–254.

Martin V, Zhao J, Afek A, Mielko Z, Gordân R. 2019. QBiC-Pred: quantitative predictions of transcription factor binding changes due to sequence variants. Nucleic Acids Research 47: W127–W135.

de Santiago I, Liu W, Yuan K, O’Reilly M, Chilamakuri CSR, Ponder BAJ, Meyer KB, Markowetz F. 2017. BaalChIP: Bayesian analysis of allele-specific transcription factor binding in cancer genomes. Genome Biol 18: 39.

Li H, Handsaker B, Wysoker A, Fennell T, Ruan J, Homer N, Marth G, Abecasis G, Durbin R, 1000 Genome Project Data Processing Subgroup. 2009. The Sequence Alignment/Map format and SAMtools. Bioinformatics 25: 2078–2079.

Bressloff PC. Stochastic processes in cell biology. Vol. 41. Berlin: Springer, 2014.

Osorio, D., Yu, X., Yu, P., Serpedin, E., and Cai, J.J. (2019). Single-cell RNA sequencing of a European and an African lymphoblastoid cell line. Sci Data 6, 112.

Rabani M, Levin JZ, Fan L, Adiconis X, Raychowdhury R, Garber M, Gnirke A, Nusbaum C, Hacohen N, Friedman N, et al. 2011. Metabolic labeling of RNA uncovers principles of RNA production and degradation dynamics in mammalian cells. Nat Biotechnol 29: 436–442.

Shahrezaei V, Swain PS. 2008. Analytical distributions for stochastic gene expression. PNAS 105: 17256– 17261.

Vu TN, Wills QF, Kalari KR, Niu N, Wang L, Rantalainen M, Pawitan Y. 2016. Beta-Poisson model for single-cell RNA-seq data analyses. Bioinformatics 32: 2128–2135.

Zoller B, Little SC, Gregor T. 2018. Diverse Spatial Expression Patterns Emerge from Unified Kinetics of Transcriptional Bursting. Cell 175: 835-847.e25.

Gao T, Qian J. 2020. EnhancerAtlas 2.0: an updated resource with enhancer annotation in 586 tissue/cell types across nine species. Nucleic Acids Research 48: D58–D64.

Davis CA, Hitz BC, Sloan CA, Chan ET, Davidson JM, Gabdank I, Hilton JA, Jain K, Baymuradov UK, Narayanan AK, et al. 2018. The Encyclopedia of DNA elements (ENCODE): data portal update. Nucleic Acids Res 46: D794–D801.

Zhou W, Machiela MJ, Freedman ND, Rothman N, Malats N, Dagnall C, Caporaso N, Teras LT, Gaudet MM, Gapstur SM, et al. 2016. Mosaic loss of chromosome Y is associated with common variation near TCL1A. Nat Genet 48: 563–568.

Hagemann-Jensen M, Ziegenhain C, Chen P, Ramsköld D, Hendriks G-J, Larsson AJM, Faridani OR, Sandberg R. 2020. Single-cell RNA counting at allele and isoform resolution using Smart-seq3. Nat Biotechnol 38: 708– 714.

Fukaya T, Lim B, Levine M. 2016. Enhancer Control of Transcriptional Bursting. Cell 166: 358–368.

Kerimov N, Hayhurst JD, Peikova K, Manning JR, Walter P, Kolberg L, Samoviča M, Sakthivel MP, Kuzmin I, Trevanion SJ, et al. 2021. A compendium of uniformly processed human gene expression and splicing quantitative trait loci. Nat Genet 53: 1290–1299.

Gertz J, Varley KE, Reddy TE, Bowling KM, Pauli F, Parker SL, Kucera KS, Willard HF, Myers RM. 2011. Analysis of DNA Methylation in a Three-Generation Family Reveals Widespread Genetic Influence on Epigenetic Regulation. PLOS Genetics 7: e1002228.

Larson DR, Zenklusen D, Wu B, Chao JA, Singer RH. 2011. Real-time observation of transcription initiation and elongation on an endogenous yeast gene. Science 332: 475–478.

George L, Indig FE, Abdelmohsen K, Gorospe M. Intracellular RNA-tracking methods. Open Biology 8: 180104.

Boegel S, Löwer M, Bukur T, Sorn P, Castle JC, Sahin U. 2018. HLA and proteasome expression body map. BMC Medical Genomics 11: 36.

Gupta A, Martin-Rufino JD, Jones TR, Subramanian V, Qiu X, Grody EI, Bloemendal A, Weng C, Niu S-Y, Min KH, et al. 2022. Inferring gene regulation from stochastic transcriptional variation across single cells at steady state. Proc Natl Acad Sci U S A 119: e2207392119.

Goes FS, McGrath J, Avramopoulos D, Wolyniec P, Pirooznia M, Ruczinski I, Nestadt G, Kenny EE, Vacic V, Peters I, et al. 2015. Genome-wide association study of schizophrenia in Ashkenazi Jews. Am J Med Genet B Neuropsychiatr Genet 168: 649–659.

Qiu P. 2020. Embracing the dropouts in single-cell RNA-seq analysis. Nat Commun 11: 1169.

Naumann, S., Reutzel, D., Speicher, M., & Decker, H. J. (2001). Complete karyotype characterization of the K562 cell line by combined application of G-banding, multiplex-fluorescence in situ hybridization, fluorescence in situ hybridization, and comparative genomic hybridization. Leukemia research, 25(4), 313–322.

Zhou, B., Ho, S. S., Greer, S. U., Spies, N., Bell, J. M., Zhang, X., Zhu, X., Arthur, J. G., Byeon, S., Pattni, R., Saha, I., Huang, Y., Song, G., Perrin, D., Wong, W. H., Ji, H. P., Abyzov, A., & Urban, A. E. (2019). Haplotype-resolved and integrated genome analysis of the cancer cell line HepG2. Nucleic acids research, 47(8), 3846– 3861.

Zhou Y, Song WM, Andhey PS, Swain A, Levy T, Miller KR, Poliani PL, Cominelli M, Grover S, Gilfillan S, et al. 2020. Human and mouse single-nucleus transcriptomics reveal TREM2-dependent and TREM2-independent cellular responses in Alzheimer’s disease. Nat Med 26: 131–142.

Bianchi A, Scherer M, Zaurin R, Quililan K, Velten L, Beekman R. 2022. scTAM-seq enables targeted highconfidence analysis of DNA methylation in single cells. Genome Biology 23: 229.

Lother A, Bergemann S, Deng L, Moser M, Bode C, Hein L. 2018. Cardiac Endothelial Cell Transcriptome. Arteriosclerosis, Thrombosis, and Vascular Biology 38: 566–574.

